# Nitrous Oxide Formation and Consumption in Thawing Permafrost: A Microcosm Study

**DOI:** 10.1101/2024.06.11.598536

**Authors:** Yanchen Sun, Xiaofen Wu, Oksana G. Zanina, Elizaveta M. Rivkina, Karen G. Lloyd, Frank E. Löffler, Tatiana A. Vishnivetskaya

**Author notes:** Corresponding authors: Tatiana A. Vishnivetskaya, University of Tennessee, Department of Microbiology, Ken and Blaire Mossman Building, 1311 Cumberland Avenue, Knoxville, TN 37996, USA. Frank E. Löffler, University of Tennessee, Department of Civil and Environmental Engineering, 325 John D. Tickle Building, 851 Neyland Drive, Knoxville, TN 37996, USA. Department of Marine Chemistry and Geochemistry, Woods Hole Oceanographic Institution, Woods Hole, Massachusetts 02543, USA. Department of Pediatrics, Washington University School of Medicine, St Louis, Missouri, USA.

## Abstract

Nitrous oxide (N_2_O) emissions contribute to stratospheric ozone depletion and global warming. Climate warming causes permafrost thawing and decomposition of the dormant nitrogenous compounds, releasing N_2_O; however, understanding of the microbial formation and consumption of N_2_O in permafrost is still limited. Permafrost soils collected at two depths (5.4 m and 16.9 m) from the East Siberian Sea coast of Russia were used to establish microcosms assessing N_2_O formation and consumption in the presence of either nitrate (NO_3_^-^, 1 mM) or N_2_O (1 mM), respectively, during incubation at 4 and 20°C. Rapid N_2_O formation was observed in NO_3_^-^-amended microcosms, but N_2_O consumption was slow and incomplete over a 1-year incubation period in all microcosms. Twenty-six quality-filtered metagenome-assembled genomes (MAGs) harboring genes involved in the reduction of NO_3_^-^ and/or N_2_O were recovered from 16 metagenomes obtained from duplicate NO_3_^-^- and N_2_O-amended microcosms. None of the MAGs carried a complete set of genes to perform canonical denitrification (i.e., NO_3_^-^→N_2_) indicating N_2_O formation and consumption is likely driven by non-denitrifying bacteria. While coastal permafrost microbiomes harbor *nosZ* genes, activity monitored in the microcosms indicates N_2_O formation exceeds N_2_O consumption, emphasizing the need for integrated approaches to assess and predict N turnover in thawing permafrost.

## Introduction

Permafrost covers approximately 17% of the global terrestrial surface [1] and preserves about 50% of the total global soil organic carbon and nitrogen [2–5]. Global warming drives the increasing thawing of permafrost, releasing a large amount of greenhouse gases (i.e., carbon dioxide [CO_2_], methane [CH_4_], and nitrous oxide [N_2_O]), exacerbating the climate-change feedback [6–10]. Under the current scenario, the temperature in Artic regions would increase by 5.6-12.4°C by the end of 2100 [11], which is predicted to lead to extensive thawing and degradation of vulnerable permafrost [12]. Presumably, thawing permafrost could be a significant source of greenhouse gases in the coming decades and centuries. The massive release of greenhouse gases may trigger yet another overlooked feedback, namely the inhibition of microbial CH_4_ production by N_2_O, which has not been considered in greenhouse gas emission models [13].

N_2_O is a pivotal greenhouse gas that has been underappreciated in traditional global greenhouse gas budgets in the Arctic [6–8, 14]. N_2_O is also a long-lived ozone-depleting gas with a global warming potential three hundred times and ten times greater than those of the equivalent amounts of CO_2_ and CH_4_, respectively [15–17]. Permafrost thaw causes the decomposition of soil organic nitrogen, ultimately leading to continued emission of N_2_O [6, 7, 14, 18]. For instance, N_2_O emissions range from up to 3 mg N_2_O m^-2^ d^-1^ from thawing permafrost peatlands [6] to about 0.9 mg N_2_O m^-2^ d^-1^ from thawing Yedoma permafrost [14], and is on average of 0.29 mg N_2_O m^-2^ d^-1^ across all types of permafrost ecosystems [7]. As a result, the observed N_2_O fluxes predict the N_2_O sources in thawing permafrost outpacing N_2_O sinks. These measurements suggest that permafrost is becoming a substantial N_2_O source. The overall emission of N_2_O is collectively determined by its formation and consumption. Therefore, detailed investigation of N_2_O formation and consumption in thawing permafrost is necessary to understand its total emissions.

Biological formation of N_2_O typically results from multiple processes, including denitrification (NO_3_^-^/NO_2_^-^→N_2_O) [19], nitrification (NH_4_^+^→NO_3_^-^) [20], dissimilatory nitrate reduction to ammonium (DNRA) (NO_3_^-^/NO_2_^-^→NH_4_^+^) [21], and chemodenitrification [22, 23]. In contrast to the diverse sources of N_2_O, the major natural biological sink of N_2_O is its reduction catalyzed by N_2_O reductase (NosZ) [24]. Studies of N_2_O emissions from permafrost and permafrost-affected soils are mainly conducted by measuring N_2_O fluxes in mesocosms or field experiments [6, 14, 18]; however, the relative contributions of N_2_O formation versus its consumption to the overall emission, as well as microorganisms involved in these two processes, have not been fully elucidated.

A recent study demonstrated that N_2_O emissions are also affected by permafrost type, with seasonally thawed active layers releasing more N_2_O than freshly thawed permafrost [14]. This observation may be due to changes in metabolism and alleviated functional limitations of microbial communities in thawed permafrost [25, 26]. However, microbial communities in the top active layer (i.e., seasonally-thawed region) could diffuse and migrate down to the deeper active layers [27] and finally accumulate on the, so called, permafrost table that prevents further penetration into perennially frozen layers, complicating the investigation of microorganisms involved in N_2_O formation and consumption. In the current study, we used permafrost soils collected from the East Siberian Sea coast of Russia, which have not experienced any documented freeze-thaw cycles [28, 29]. Furthermore, comparative analysis of the pristine permafrost metagenomes revealed nitrogen metabolic potential [29], which warrants our detailed investigation of microbial N_2_O formation and consumption.

The objectives of this study were to examine N_2_O formation and consumption in thawing permafrost soil microcosms and identify the key microorganisms driving these two contrasting processes. To achieve these objectives, we explored, in a series of microcosms, the reduction of NO_3_^-^ and N_2_O following a 1-year long incubation under environmentally relevant temperature conditions (4°C versus 20°C) [8, 14, 30], and monitored responses of the microbial community through N_2_O measurements and analyses of metagenomes.

## Materials and Methods

### Permafrost samples information

Permafrost samples (formed about 100-120 kyr ago) were collected from the East Siberian Sea coast, the northern boundary of the Kolyma Lowland in northeastern Siberia, Russia, in August 2017 at depths of 5.4 and 16.9 m in the same borehole (Figure S1) as described [29]. Core samples were treated with an aseptic sampling protocol [31]. The silt loams from the upper horizon (depth of 5.4 m) of the borehole were identified to be coastal brackish permafrost (defined as 54BP) and deeper layers (depth of 16.9 m) were recognized to be saline silt loams and sandy loams of marine permafrost (defined as 169MP). Different nitrogen species (e.g., nitrate, nitrite, ammonium) were detected in the coastal permafrost deposits [32]. The physicochemical properties of the permafrost samples were analyzed and described previously (Table S1) [29].

### Permafrost microcosms

Bicarbonate-buffered mineral salts medium was prepared following established protocols [33]. L-cysteine (0.2 mM) was added as a reductant. Microcosms were set up according to an established procedure [34]. Briefly, about 8 g of each pristine permafrost sample, 54BP and 169MP, were homogenized aseptically and 1 g of permafrost material was transferred to 60-mL glass serum bottles containing 30 mL of medium using sterilized stainless-steel spatulas inside a glove box (Coy Laboratory Products, Grass Lake, MI, USA) filled with 97% N_2_ and 3% H_2_. The serum bottles were immediately sealed with sterile butyl rubber stoppers, crimped with aluminum caps, and removed from the glove box. Pyruvate (5 mM) was added to each serum bottle from an anoxic, filter-sterilized 1 M stock solution by syringe and replenished after 6 and 12 months. A total of 0.3 mmol of pyruvate was added to each microcosm over 1-year of incubation. In addition, the Wolin vitamin mix and CuCl_2_ (∼17 μM) were added from concentrated stock solutions to individual bottles [35]. Each setup had two replicated microcosms (Figure S1) due to the limited amounts of permafrost soil available. Plastic syringes with 25-gauge needles (BD, Franklin Lakes, NJ, USA) were used to add 1 mM NO_3_^-^ (30 µmol) from a 1 M stock solution or 1 mM N_2_O (2 mL or 80 µmol of undiluted N_2_O). N_2_O was analyzed every 20-30 days across the incubation using gas chromatography (GC) (see below) [36]. NO_3_^-^ and NO_2_^-^ were periodically analyzed using ion chromatography (IC) by extracting liquid samples from the microcosms [23]. The bottles were incubated at 4 and 20°C under static conditions. Negative controls included heat-killed (autoclaved) replicates and microcosms without N_2_O and NO_3_^-^ but with pyruvate at pH 7.2 for each permafrost sample.

### Analytical procedures

N_2_O was analyzed with an Agilent 3000A Micro-GC (Agilent, CA, USA) equipped with a thermal conductivity detector and a Plot Q column [34, 37]. The injector and column temperatures were set to 100°C and 50°C, respectively, and the column pressure was set to 25 psi. Gas sample (0.1 mL) was withdrawn from the microcosm headspace and manually injected into the Micro-GC. Aqueous N_2_O concentrations were calculated from the headspace concentration using a dimensionless Henry’s constant for N_2_O based on the equation *C*_aq_ = *C*_g_/*H*_cc_ [38]. *C*_aq_, *C*_g_, and *H*_cc_ are the aqueous N_2_O concentration (μM), the headspace N_2_O concentration (μM), and the Henry’s constant (dimensionless), respectively. The total amount of N_2_O equaled the sum of N_2_O in the headspace and aqueous phase.

NO_3_^-^ and NO_2_^-^ were quantified by ion chromatography using a reagent-free eluent regeneration system (ICS-2100; Dionex, Sunnyvale, CA) and a Dionex IonPac AS18 4- by 250- mm analytical column heated to 30°C. The eluent was 10 mM KOH and the flow rate was 1 mL min^-1^. For each measurement, a 0.5 mL liquid sample was withdrawn from the microcosm and filtered through the 0.2 μm PES filter (Nalgene, Rochester, NY, USA), and then the NO_3_^-^ and NO_2_^-^ were measured in the filtrate.

### DNA extraction and metagenome sequencing

Suspension aliquots (5 mL) were sub- sampled from the homogenized microcosms with 5 mL plastic syringes equipped with 18-gauge needles after 1-year incubation. The total genomic DNA for untargeted shotgun metagenome sequencing was extracted with the DNeasy PowerSoil kit (Qiagen, Hilden, Germany) following the manufacturer’s protocol. DNA concentrations were determined using the Qubit fluorometer (Life Technologies, Carlsbad, CA). Sequencing of the metagenomic DNA was performed at the Genomics Core at the University of Tennessee using the NovaSeq 6000 platform (NovaSeq 6000 SP reagent kit, Illumina, San Diego, CA) to generate metagenomic datasets with 2 × 250 bp read length (Table S2).

### Bioinformatic analyses

The raw metagenomic reads derived from two original permafrost samples (Table S3, PRJNA601698) [29] and 16 samples obtained from NO_3_^-^- and N_2_O-amended microcosms were trimmed using Trimmomatic v.0.39 with default settings [39]. Assembly of the trimmed short reads was performed using MEGAHIT v.1.2.9 [40], and contigs longer than 1,000 bp in length were routed into downstream analyses. Metagenomic data from two replicates of the same type of microcosm were co-assembled. Contigs were binned using MaxBin2 v.2.2.4 [41], MetaBAT v.2.12.1 [42], and CONCOCT v.1.1.0 [43] with default settings, respectively, to recover individual metagenome-assembled genomes (MAGs). An optimized, non-redundant set of MAGs was recovered using metaWRAP with the command bin_refinement from MAGs generated by using three different binning packages [44]. The resulting MAGs were checked for completeness and contamination using CheckM v.1.0.18 [45]. The coverage of each MAG was calculated after removing bases with the highest and lowest 5% coverage by using CoverM v.0.6.1 (https://github.com/wwood/CoverM). The relative MAG abundance in each sample was defined as the number of reads aligning to each contig of the MAG normalized by the total number of reads in the sample [46].

### Microbial community profiling

GraftM v.0.13.1 was used to extract 16S rRNA gene fragments from the trimmed metagenomic datasets and those 16S rRNA gene sequences were classified using the default Greengenes database (release 13_8) at the 97% nucleotide identity level [47, 48]. The relative abundance of the operational taxonomic units (OTUs) was calculated based on the number of reads assigned to each OTU. OTUs were used instead of amplicon sequence variants (ASVs) because, as the data come from metagenomes and not primer-amplified genes, individual sequences vary slightly in length. The microbial community profile was assessed based on OTU taxonomic assignments at the phylum and family levels.

### Taxonomy, functional annotation, and comparative genomics

MAGs with completeness >50% and contamination <10% were designated as quality-filtered MAGs, which were routed into subsequent downstream analysis. Taxonomic assignments for each quality-filtered MAG were conducted by using GTDB-Tk v.0.1.4 [49] based on the Genome Taxonomy Database (GTDB, http://gtdb.ec/ogenomic.org) taxonomy RS207 [50]. Protein-coding sequences present in quality-filtered MAGs were predicted using Prodigal v.2.6.3 [51], and assigned Kyoto Encyclopedia of Genes and Genomes (KEGG) orthologs (KOs) using KofamScan v.1.3.0 against hidden Markov model (HMM) profiles from the KEGG database (released on Sep-20-2022) [52]. The completeness of various metabolic pathways was assessed using KEGG-Decoder v.1.32.0 [53], and pathways of interest (e.g., nitrogen cycling, lactate, and hydrogen metabolism, copper transport) were specifically selected from the assessed results. Average amino acid identity (AAI) values between the quality-filtered MAGs from the current study and close relatives from formally named microbial species (NCBI Genome database) were calculated using the Microbial Genome Atlas (MiGA) (http://microbial-genomes.org/). MAGs encoding N-cycling genes were collected based on the KEGG annotation results. The phylogenetic tree of quality-filtered MAGs was created using FastTree v.2.1.8 (WAG+GAMMA models) with a concatenated alignment of 120 bacterial and 122 archaeal conserved marker genes [54]. The generated tree files were visualized with iTOL [55].

### Statistical analyses

Statistical analyses were performed using R v.4.0.2 [56]. Beta-diversity was calculated using weighted-UniFrac distances and visualized using the principal coordinate analysis (PCoA) plot in R with packages ggplot2 [57] and phyloseq [58]. Statistical differences in microbial communities among original permafrost soils, microcosms at 4°C, and microcosms at 20°C were determined using permutational multivariate analysis of variance (PERMANOVA) using the adonis function in vegan with 999 permutations [59]. KEGG annotation results of quality-filtered MAGs were shown using heatmap in R with package pheatmap [60].

## Results

### N_2_O formation and consumption in permafrost soil microcosms

NO_3_^-^ reduction from 30 µmol to 20 µmol and almost to 0 µmol occurred in microcosms with 54BP permafrost soil collected from 5.4 m depth at both 4°C and 20°C (Figure S2A); however, NO_3_^-^ reduction from 30 µmol to 10 µmol was only observed in one microcosm at 4°C with 169MP permafrost soil from 16.9 m depth (Figure S2B). After a 1-year, microcosms established with soils from both depths and incubated at 4°C yielded NO_2_^-^ and N_2_O (Figures S2C and 1A). N_2_O generated during NO_3_^-^ reduction plateaued after 3 months of incubation, and up to 3.2 and 2.2 μmol of N_2_O were measured in the NO_3_^-^-amended microcosms using 54BP and 169MP permafrost samples, respectively (Figure 1A). In contrast, NO_3_^-^ reduction at 20°C in the microcosm using 54BP permafrost transiently produced 6.3 μmol NO_2_^-^ (Figure S2D) and the final product was N_2_O up to 11.5 μmol (Figure 1B). No N_2_O was detected in control microcosms that did not receive NO_3_^-^, indicating that N_2_O formation from nitrogenous compounds in the medium or associated with the permafrost soil did not occur or was negligible.

**Figure 1.**
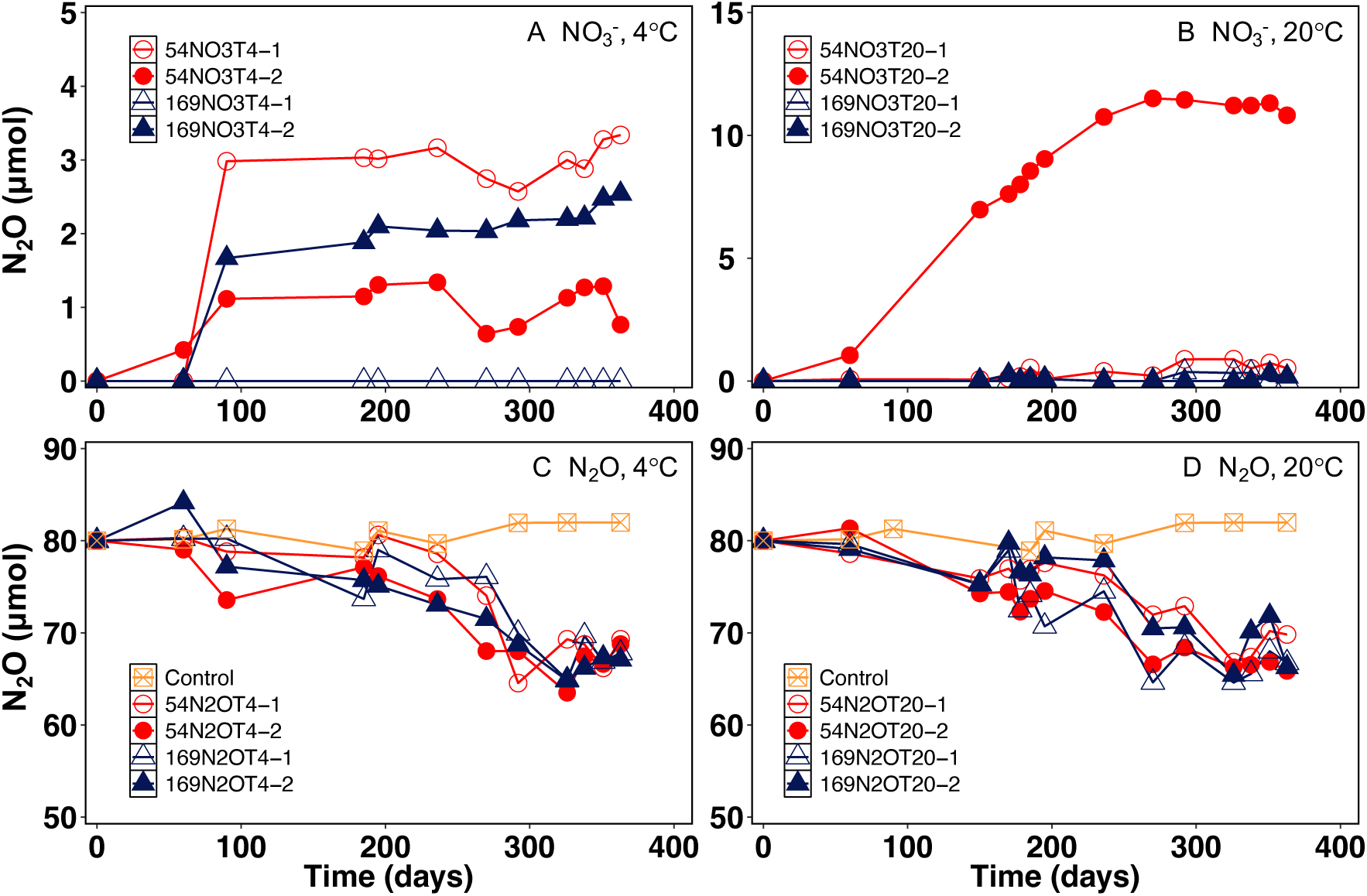
N_2_O profiles in permafrost soil microcosms. Panels A and B show N_2_O production in NO_3_^-^-amended microcosms stimulated by incubation at 4°C and 20°C, respectively. Panels C and D show the N_2_O consumption in N_2_O-amended microcosms at 4°C and 20°C, respectively. The red circles and blue triangles represent the permafrost samples from depths of 5.4 m and 16.9 m, respectively. The orange squares in panels C and D represent the heat-killed control. Each type of microcosm has two replicates shown as open symbols for replicate 1 and closed symbols for replicate 2. Heat-killed control has one replicate.

Measurable N_2_O consumption occurred in all N_2_O-amended microcosms at 4°C (Figure 1C) and 20°C (Figure 1D). Over the course of the 1-year incubation period, the average N_2_O consumption (n=2) in the 5.4 m and 16.9 m microcosms at 4°C was 10.9 and 12.6 μmol of N_2_O, respectively, with an average reduction rate of 0.032 μmol day^-1^. At 20°C, 12.2 and 13.3 μmol of N_2_O were reduced in 5.4 m and 16.9 m microcosms, respectively, after a 1-year incubation, with an average reduction rate of 0.035 μmol day^-1^. Evidently, thawing permafrost soils have the potential for biological reduction of N_2_O.

### Microbial community composition

At the end of the incubation period, the microcosms were sacrificed and the total genomic DNA was extracted from 5 ml of slurry. The 16 metagenomic datasets generated from the individual microcosms (Table S2), as well as three datasets generated from the original permafrost soils (Table S3), revealed a total of 1,055 16S-rRNA gene-based OTUs (Table S4), and the average number of OTUs detected in microcosms amended with either N_2_O or NO_3_^-^ was 1.2-3.4 fold lower than in the corresponding pristine permafrost soils. One exception was the NO_3_^-^-amended microcosms for the 54BP sample at 20°C where the average number of OTUs was 1.2 fold higher than in pristine 5.4 m permafrost soil (Table S4). Beta diversity analysis, a measure of the dissimilarity between communities, indicated distinct community compositions in response to depth and temperature (*p* < 0.05, PERMANOVA) (Figure 2A). PCoA showed that approximately 63% of the total variability of the microbial communities at the OTU level is explained by depth and incubation temperature (Figure 2A and Table S5). The analysis shows that the communities from pristine permafrost samples 54BP and 169MP group together but the microcosm enrichment had a profound impact on the microbiomes. The communities that developed at 4°C in NO_3_^-^-amended microcosms irrespective of depth grouped closely together. The 4°C communities from 169MP N_2_O-amended microcosms group separately but close to either communities from original 169MP permafrost or 169MP microcosms incubated at 20°C regardless of substrate (Figure 2A). The communities that developed at 20°C in both 169MP NO_3_^-^- and N_2_O-amended microcosms are most similar to each other and group separately from the rest of the microcosms and the two original permafrost soils. The community composition of 54BP NO_3_^-^- and N_2_O-amended microcosms incubated at 20°C show high variations between replicates with one 54BP N_2_O replicate incubated at 20°C showing a similar microbial profile to the 54BP microcosms incubated at 4°C (Figure 2A).

**Figure 2.**
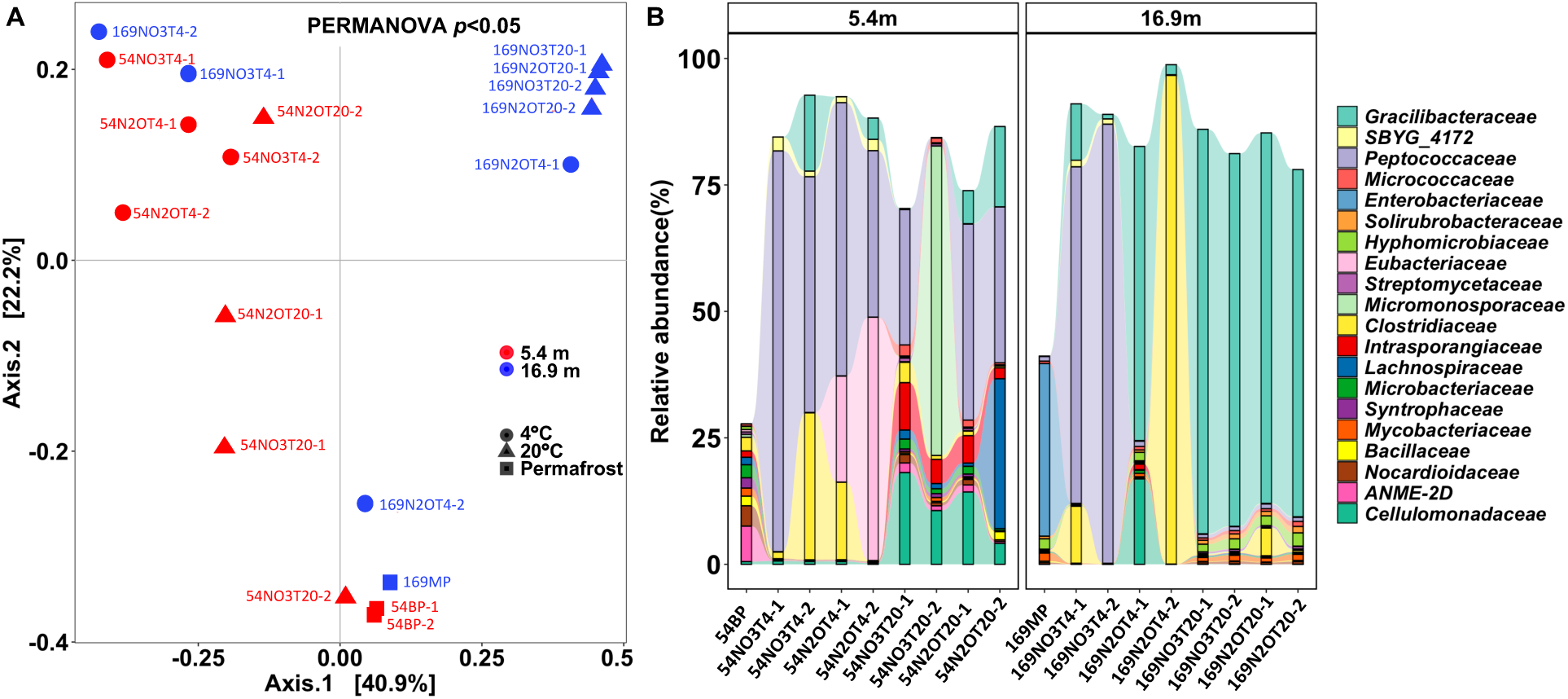
Microbial community composition identified in the original permafrost, NO_3_^-^-, and N_2_O-amended microcosms incubated at 4°C and 20°C based on 16S rRNA gene fragments recovered from the metagenomes. (A) Beta diversity of microbial communities based on weighted UniFrac analysis of 16S rRNA gene fragments recovered from the metagenomes. Relationship between samples were visualized by principal coordinates analysis (PCoA): colors distinguish the depth where samples were collected; shape indicates the original permafrost sample or microcosms at 4°C and 20°C; source of nitrogen and replica are indicated in the name. (B) The relative abundance distributions of the top 20 families observed in the original permafrost and the different microcosms. The x-axis labels indicate individual samples, except 54BP where the average community is presented. Sample names contain reference to depth, source of nitrogen, incubation temperature (T4 means 4°C and T20 means 20°C), and replicate.

The majority of the 16S rRNA gene sequences from the pristine 54BP permafrost are affiliated with four major phyla namely *Actinobacteria*, *Euryarchaeota*, *Firmicutes*, and *Proteobacteria*, with a combined relative abundance exceeding 80% (Figure S3). *Proteobacteria*, *Chloroflexi*, *Actinobacteria*, and *Crenarchaeota* dominate the pristine 169MP permafrost with a combined relative abundance of 87% (Figure S3). In contrast, the NO_3_^-^ and N_2_O-amended microcosms at 4°C are dominated by *Firmicutes* (except one replicate (169N2OT4-1) for the 16.9 m N_2_O condition where ∼30% of *Actinobacteria* was observed), while microcosms at 20°C contain both Firmicutes (15-90%) and *Actinobacteria* (10-18%) (Figure S3).

Community analysis showed that sequences affiliated with *Peptococcaceae*, *Gracilibacteraceae*, and *Clostridiaceae* families of the phylum *Firmicutes* increased in relative abundance in NO_3_^-^-amended microcosms using both 54BP and 169MP permafrost at 4°C (Figure 2B). In contrast, the 54BP NO_3_^-^-amended microcosms incubated at 20°C are dominated by *Cellulomonadaceae* (*Actinobacteria*) sequences and either *Peptococcaceae* (*Firmicutes*) or *Micromonosporaceae* (*Actinobacteria*), while microbial communities in 169MP microcosms at similar conditions are dominated by *Gracilibacteraceae* (*Firmicutes*). In the case of N_2_O- amended microcosms incubated at 4°C, *Peptococcaceae* and *Eubacteriaceae*, both in the phylum *Firmicutes*, dominate the 54MP microcosms, whereas *Gracilibacteraceae* and *Clostridiaceae*, also members of the phylum Firmicutes, dominate the 169MP microcosms. The N_2_O-amended microcosms at 20°C yielded a more diverse community in 54BP microcosms where *Gracilibacteraceae*, *Peptococcaceae*, *Lachnospiraceae* families from the phylum *Firmicutes* and *Cellulomonadaceae* family from the phylum *Actinobacteria* dominate, while the family *Gracilibacteraceae* (*Firmicutes*) dominates the 169MP microcosms (Figure 2B).

### MAGs harboring denitrification and/or DNRA genes

Among 104 quality-filtered MAGs recovered from the 16 metagenomes generated from the NO_3_^-^- and N_2_O-amended microcosms, 26 MAGs harbor genes predicted to be involved in denitrification and/or DNRA (Figure 3, Tables S6 and S7). These 26 MAGs are assigned to five phyla, with *Actinobacteria* (14 MAGs) being the most abundant and followed by *Firmicutes* (8 MAGs), *Chloroflexota* (2 MAGs), *Proteobacteria* class *Alphaproteobacteria* (1 MAG), and *Thermoproteota* (1 MAG). Most of these MAGs are assigned to novel taxa according to the AAI results (Table S8), indicating an unexplored microbial diversity related to N_2_O formation and consumption in these permafrost soils. Eight of these 26 MAGs account for 2.1-48.1% of the corresponding metagenome they were assembled from, whereas the relative abundance of the remaining 18 MAGs is below 2% (Figure 4). Of note, two MAGs affiliated with the phylum Firmicutes, 169NO3T4-bin.5 and 169NO3T4-bin.8, derived from 169MP NO_3_^-^-amended 4°C microcosms were the highest in relative abundance, reaching approximately 48% and 15% (Figure 4), respectively. Most MAGs exhibit lower or negligible relative abundances in the metagenomes derived from the original permafrost soils than in metagenomes from the matched microcosms (Figure S4), suggesting that thawing and incubation under different conditions selects for distinct microbes.

**Figure 3.**
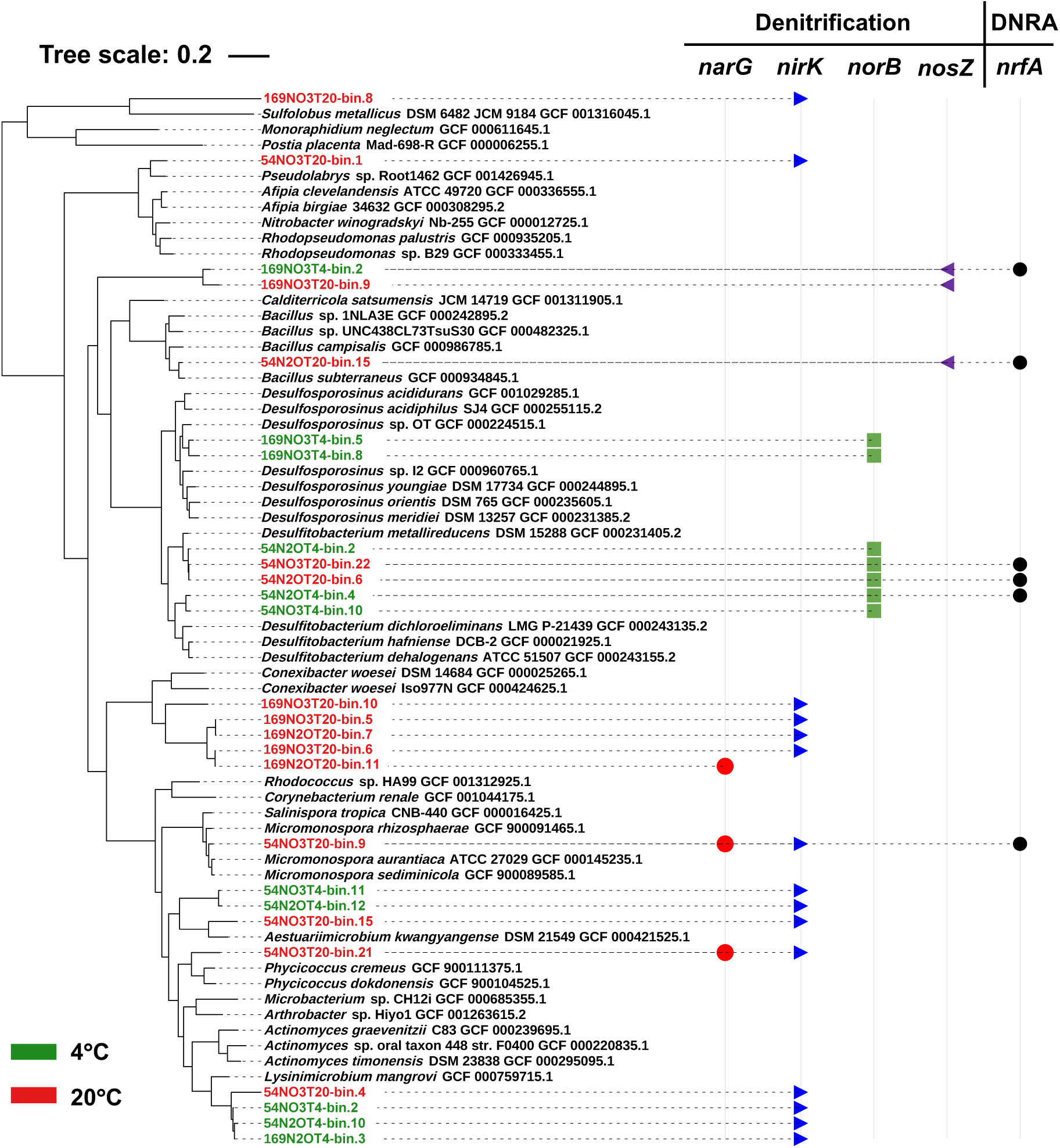
Phylogenetic analysis of MAGs harboring genes for denitrification and/or DNRA. The tree shows a phylogenetic relationship of the 26 MAGs to their closest neighbors and what genes for denitrification and/or DNRA they harbor. MAGs recovered from 4°C and 20°C microcosms are shown in green and red font, respectively. The MAGs’ names show depth, source of nitrogen, incubation temperature (T4 [4°C] and T20 [20°C]), and bin number.

**Figure 4.**
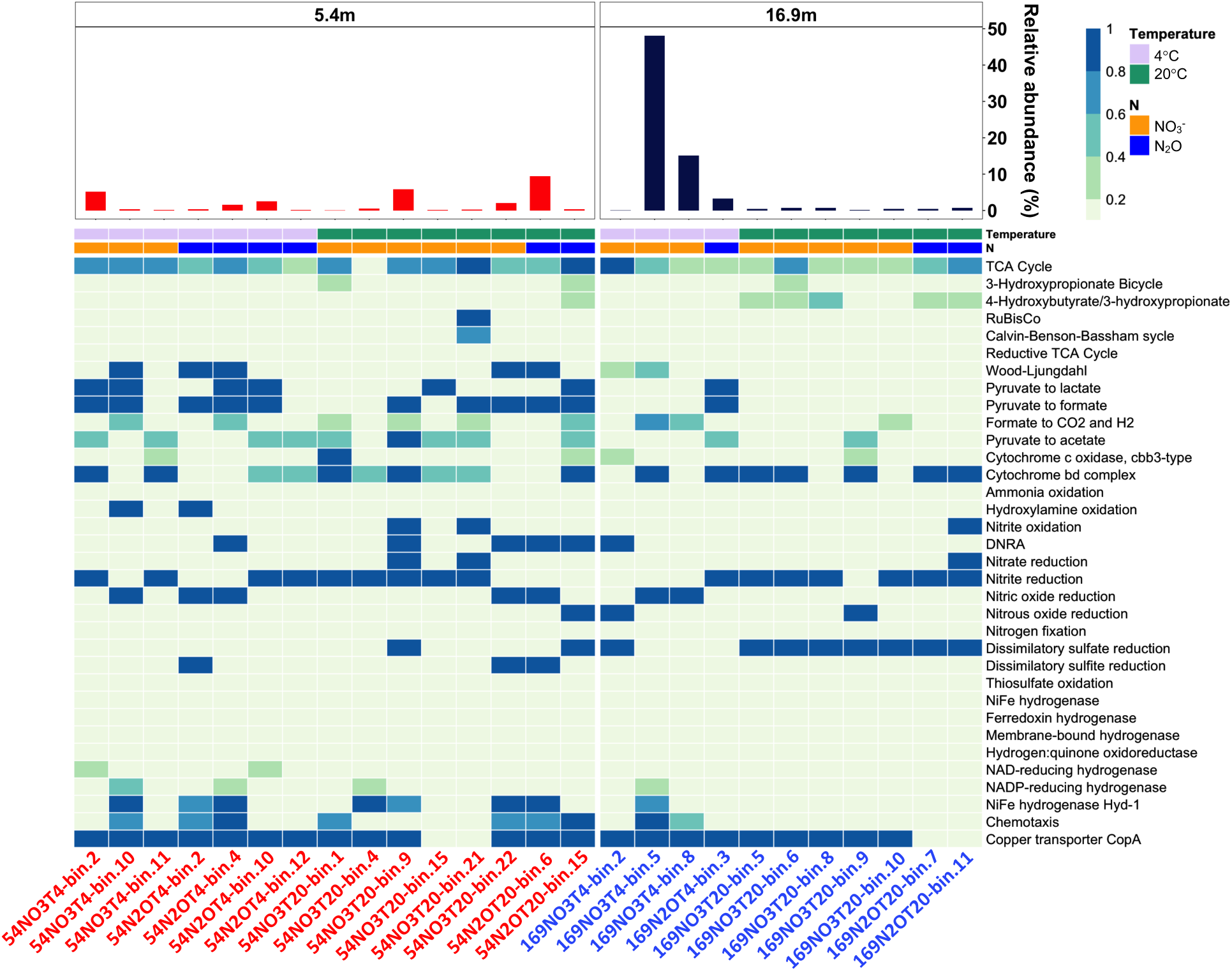
The relative abundance of MAGs harboring N cycling genes derived from the corresponding microcosms (upper panel). Functional analysis of MAGs harboring N-cycling genes (lower panel). The heatmap shows the completeness of key metabolic pathways or functions in the 26 MAGs harboring N-cycling genes based on KEGG annotation. MAGs recovered from microcosms using permafrost from 5.4 m and 16.9 m are shown in red and blue, respectively. The MAG names shown on the x-axis indicate the sample, substrate, and incubation temperatures (T4 [4°C] and T20 [20°C]).

The metagenomes generated from two original permafrost samples yielded 14 quality-filtered MAGs, namely two MAGs from 54MP and 12 MAGs from 169MP, that encoded genes for denitrification and/or DNRA and belonged to six phyla including *Thermodesulfobacteriota* (4 MAGs), *Proteobacteria* (class *Gammaproteobacteria*) (3 MAGs), *Mixococcota* (1 MAG), *Gemmatimonadota* (1 MAG), *Actinomycetota* (2 MAGs), and *Chloroflexota* (3 MAGs) (Figure S5 and Table S9). These 14 MAGs have a relative abundance ranging from 0.1% to 3.6% as estimated by mapping the trimmed reads to the corresponding MAGs (Figure S6).

### Functional analysis of MAGs harboring genes for denitrification and/or DNRA

Key metabolic pathways or functions of the MAGs harboring denitrification and/or DNRA genes were predicted based on KEGG annotation and shown in Figures 3 and 4 for 26 MAGs from microcosms and in Figures S5 and S6 for 14 MAGs from pristine permafrost. None of the MAGs from microcosms carry a complete set of genes to perform canonical denitrification (i.e., NO_3_^-^ →NO_2_^-^→ NO→N_2_O→N_2_), and only one MAG (54NO3T20-bin.9) related to the genus *Micromonospora*, phylum *Actinobacteria*, encoded the whole gene set for DNRA (i.e., NO_3_^-^→NO_2_^-^→NH_4_^+^). MAGs harboring genes encoding NO_3_^-^ reductase (*narG* and/or *napA*) were identified from the pristine permafrost (4 MAGs) and microcosms at 20°C (3 MAGs). From these 7 MAGs, only 3 MAGs were from the 54BP permafrost sample and corresponding 20°C NO_3_^-^-amended microcosms and these MAGs were affiliated with *Actinobacteria* (Figure 4 and S5). Other NO_3_^-^ reductase containing MAGs were from 169MP permafrost (3 MAGs affiliated with phyla *Thermodesulfobacteriota*, *Mixococcota*, *Gemmatimonadota*) and corresponding 20°C NO_3_^-^-amended microcosms (1 *Actinobacteria* MAG). Sixteen microcosm MAGs and 4 permafrost MAGs harbor the NO_2_^-^ reduction gene *nirK* (k00368, encoding the copper-dependent nitrite reductase), but gene *nirS* (k15864, encoding the cytochrome *cd_1_* dependent nitrite reductase NirS) was not found. *nor*B genes encoding nitric oxide reductase were found in 7 microcosm MAGs, which are affiliated with the genera *Desulfosporosinus* and *Desulfitobacterium*, both strictly anaerobic bacteria belonging to the phylum *Firmicutes*. The rest of the genes responsible for denitrification were not identified in these 7 MAGs; however, 3 of these 7 MAGs contained *nrf*A genes implicated in DNRA. Only 3 MAGs from 169MP permafrost (affiliated with *Chloroflexota*, *Mixococcota*, *Alphaproteobacteria*) and 3 MAGs enriched in the microcosms, namely 54N2OT20-bin.15, affiliated with the genus *Bacillus* (phylum *Firmicutes*), and 169NO3T4-bin.2 and 169NO3T20-bin.9 affiliated with the class *Anaerolineae* (phylum *Chloroflexota*), harbor *nosZ* genes for N_2_O reduction. No other genes related to denitrification were identified in these MAGs, but only 1 MAG from the original 169MP and 2 microcosms’ MAGs carry gene *nrf*A for DNRA.

In addition to the permafrost associated organic substrates (Table S1), pyruvate was used as the carbon source and electron donor for the reduction of NO_3_^-^ and N_2_O. About 43% of permafrost MAGs and 46% of microcosms’ MAGs harboring denitrification and/or DNRA genes also carry genes for metabolizing pyruvate to acetate, formate, and lactate (Figure 4 and S6). About 86% of permafrost MAGs show potential for metabolizing pyruvate (Figure S6). These findings indicate that the majority of MAGs harboring denitrification and/or DNRA genes are capable of directly utilizing pyruvate as an electron donor. The last step of the denitrification process, the reduction of N_2_O to N_2_, is catalyzed by the well-characterized copper-containing enzyme *nos*Z and the expression of *nosZ* is controlled by the extracellular copper concentration [61]. Genes for copper transport were identified in almost 65% of the permafrost MAGs and 85% of the microcosms’ MAGs, indicating that most of the MAGs harboring denitrification and/or DNRA genes capable of taking up copper from the environment.

## Discussion

### Current paradigms for studying N_2_O emissions from permafrost

Global warming promotes permafrost degradation, replenishing the inorganic N pool (i.e., NH_4_^+^ and NO_3_^-^) through the mineralization of organic compounds and nitrification and denitrification [62, 63]. Transformation of these two N species (NH_4_^+^ and NO_3_^-^) can lead to the formation of N_2_O, as shown by our microcosm results. Though N_2_O emissions from permafrost-affected areas, particularly from active layers that experience seasonal freeze-thaw cycles, have been investigated in both field and microcosms settings [6–8, 14, 18], knowledge about microbes implicated in N_2_O formation and consumption is scarce. In addition, microbial communities in the thawing permafrost can rapidly change metabolism [25, 26]. Permafrost or perennially frozen soils preclude any diffusion and microbial dispersal down from the active layer, as well as lateral and vertical migration through the permafrost. Even though the molecular diameter of gasses including N_2_O (3.32 × 10^-10^ m) is smaller than the thickness of unfrozen water films in permafrost (5 × 10^-10^ m), diffusion of the gasses, including N_2_O, at temperatures below freezing is negligible [25]. Given the wide range of projections for permafrost thaw in the coming decades [11, 64], studies investigating N_2_O emissions using undisturbed permafrost contribute to the understanding of future N_2_O emissions following permafrost thaw.

### Thawing of permafrost yields N_2_O

The experimental results from the current study reveal that N_2_O can be slowly reduced in thawed permafrost microcosms; however, a positive N_2_O flux was monitored in the NO_3_^-^-amended microcosms under the same conditions (Figure 1). Of note, the formation of N_2_O indicates that the thawing permafrost could be an underappreciated source of N_2_O from incomplete denitrification. Our findings are consistent with N_2_O emissions observed in field and microcosm studies [6, 14, 65] and further underscore that permafrost is a major and increasingly recognized source of N_2_O [7]. The microbial communities inhabiting the permafrost soils in the current study are capable of generating N_2_O from NO_3_^-^, indicating that the permafrost harbors the metabolic potential to generate N_2_O, a finding a permafrost microcosm study corroborates [14]. The samples used in the current study are perennially frozen brackish and marine sediments from the East Siberian Sea coast and these sediments are distributed along the coast of the Arctic seas and the Arctic Ocean in Russia, Alaska, and Canada [66]. Coastal permafrost environments may become important sources of N_2_O during changes due to climate warming.

Denitrification is one of the major processes releasing N_2_O from thawed permafrost [6, 65, 67, 68]. The thawing of permafrost promotes the decomposition of soil organic matter, replenishes the NO_3_^-^/NO_2_^-^ pools, and further fuels denitrification [6]. Our study demonstrates that the reduction of NO_3_^-^ stalled at the end product of N_2_O (up to 11 μmol) and negligible N_2_O reduction was observed after the complete reduction of NO_3_^-^ (Figures 1A and 1B), emphasizing that thawing permafrost could be a major source of N_2_O [14]. In contrast, slow N_2_O reduction occurred in N_2_O-amended permafrost microcosms (Figures 1C and 1D). Collectively, the findings show that different nitrogen species (e.g., NO_3_^-^, NO_2_^-^, NH_4_^+^) preserved in coastal permafrost will contribute to a NO_3_^-^ pool in thawing permafrost and promote the emission of N_2_O via incomplete denitrification. Permafrost-affected areas undergoing episodic seasonal warming and cooling are considered major N_2_O sources [14], and the capability of N_2_O reduction suggests that in situ N_2_O consumption may occur over the longer term, raising questions about the impact of environmental change on N_2_O fluxes from this relevant ecosystem.

### Non-denitrifiers responsible for N_2_O formation and consumption

Multiple known N cycling processes, including denitrification, nitrification, DNRA, and chemodenitrification contribute to N_2_O formation in natural environmental systems [69, 70]. However, N_2_O generated in the NO_3_^-^-amended thawing permafrost microcosms may experience a continuous reduction process driven by multiple non-denitrifying bacteria due to none of these MAGs harbor the complete gene set for reducing NO_3_^-^ to N_2_O. The comparative analysis of MAGs derived from the NO_3_^-^-amended microcosms revealed that genes encoding NO_3_^-^ reductase are only in MAGs affiliated with families of *Micromonosporaceae* and *Dermatophilaceae*, genes for NO_2_^-^ reduction are mainly encoded by members of the families *Demequinaceae* and *Micromonosporaceae*, and the NO reductase gene is predominately carried by *Desulfitobacteriaceae* (Figures 3 and 4). In addition, none of these MAGs harbor the complete gene set for reducing NO_3_^-^ to N_2_O; however, the reduction of NO_3_^-^ yields NO_2_^-^ and is subsequently reduced to N_2_O (Figures 1A, 1B, S2C, and S2D). Detailed analyses of MAGs obtained from 5.4 and 16.9 m samples of original permafrost soils performed in the previous study [29] did not identify any MAGs harboring the complete gene sets for denitrification (Figures S5 and S6), which is consistent with the current findings in the NO_3_^-^-amended microcosms. We considered that the analysis of MAGs missed functional genes; however, most MAGs are of high quality (completeness >75% and contamination <5%), and at this level of completeness, genes not assembled in a MAG likely represent mobile and hypothetical genes, rather than characterized functional genes [71]. These findings reveal that multiple incomplete denitrifiers, such as *Micromonosporaceae*, *Demequinaceae*, and *Desulfitobacteriaceae*, synergistically contribute to the N_2_O formation in the permafrost soils investigated in the current study.

The only known biological N_2_O reduction process is driven by microbial N_2_O reductase (NosZ), Clade I or Clade II NosZ [69, 70]. None of the MAGs harboring *nosZ* genes and derived from either the microcosms or the original permafrost soils encode any other denitrification genes (Figures 3, 4, and S6). Furthermore, the phylogenetic analysis of the *nosZ* genes derived from the microcosms reveals that all affiliate with Clade II type *nosZ* (Figure S7), which are typically encoded by non-denitrifiers [69, 70, 72]. Collectively, our study suggests that N_2_O reduction in the microcosms is driven by non-denitrifying N_2_O reducers.

The discussion above offers a sensible explanation regarding non-denitrifying bacteria responsible for NO_3_^-^ and N_2_O reduction. Our work focuses on identifying the NO_3_^-^- and N_2_O- reducing taxa in thawing permafrost; however, replicate microcosms established with homogenized materials perform inconsistently. This variability has been reported in similar soil microcosm studies [34], but the reason for this variability remains debated [73]. The primary goal of our study is to assess whether enrichment conditions select for specific NO_3_^-^ and N_2_O- reducing taxa, and our current experimental approach achieves this goal by discovering non-denitrifying bacteria involved in the formation and consumption of N_2_O in permafrost microcosms.

### Effects of permafrost type and microcosm temperature on N_2_O emissions

A striking performance difference in microcosms using the 54BP and 169MP permafrost samples was that NO_3_^-^ was effectively reduced in all 54BP microcosms under both temperature conditions, but was reduction was slow or negligible in the 169MP microcosms irrespectively of temperature. The difference in NO_3_^-^ reduction could be due to differences in microbial abundance and diversity in the permafrost soils [29] due to different physicochemical properties and salinity (Table S1). Previous studies provide evidence that the origin, age, and physicochemical properties of permafrost determine the number of microorganisms and their taxonomic diversity [25, 74, 75]. The 16.9 m deep permafrost used in the current study was formed ∼105-120 kyr ago and is about 20 kyr older than permafrost collected at 5.4 m [25], exhibiting two times lower total carbon and total nitrogen content (Table S1) [29]. It has been reported that microbial abundance is negatively correlated with the permafrost age and positively correlated with the carbon content [25, 76]. Therefore, the microbial community associated with the deeper permafrost may have lost metabolic capabilities, including the ability to reduce NO_3_^-^ and/or metabolize carbon substrates (i.e., pyruvate).

Another plausible explanation for different NO_3_^-^ reduction results in microcosms using the 54BP and 169MP permafrost samples is the different salinity of the original permafrost. The salinity of the 169MP sample collected from the 16.9 m deep permafrost layer is 5.6 ppt (12 g L^-1^) and this is 3 times higher than that of the 54BP sample from 5.4 m permafrost (1.3 ppt or 2.8 g L^-1^). Concentrations of divalent ions including CO_3_^2-^, Ca^2+^, and Mg^2+^ were not significantly different in 54BP and 169MP samples whereas concentrations of SO_4_^2-^ and monovalent ions (Cl^-^, K^+^, and Na^+^) were significantly higher in saline 169MP (Table S1). Previous experiments with an anaerobic denitrifying microbial community showed that the NO_3_^-^ and NO_2_^-^ reduction rates are inhibited with increasing NaCl concentrations and the denitrification process in general was affected by the total ionic strength of the samples [77]. The formation of unfrozen thin brine films in permafrost provides microhabitats for microorganisms where microbial communities interact, evolve, and adapt in response to selective pressure present in native ecological niches [78, 79]. As a result, high salinity affected microbial community composition resulting in the selection of salt-tolerant microbes with lower NO_3_^-^ reduction capability and as a result hindered NO_3_^-^ reduction [80, 81].

Temperatures in the active layer of permafrost can reach as high as 20°C during the summer season [8, 14, 30]. Upon thawing, permafrost microbial communities will be impacted by rising temperatures, which may lead to rapid changes in the metabolic potential of microbial populations [26]. Therefore, incubation temperature may also be a controlling factor for differences in NO_3_^-^ reduction and N_2_O formation activity. The permafrost microbial community is viewed as the result of a continuous selection of microorganisms adapted to permafrost conditions and capable of geologically long-term coping with cold stress [26]. In our study, the incubation temperature of permafrost microcosms (i.e., 4 and 20°C) exceeds the average *in situ* temperature of permafrost (i.e., -8°C), potentially impacting the NO_3_^-^ reduction capability of cold-adapted microorganisms [82, 83].

### Environmental implications

Global warming continues to intensify, and by 2100, approximately 50% of permafrost is predicted to be lost [84–86], which will lead to continued emission of N_2_O [6, 7, 14, 18]. Understanding N_2_O formation and consumption, as well as the microorganisms involved in N turnover, is, therefore, crucial for deciphering N_2_O emissions from permafrost and predicting the feedback of N_2_O fluxes to climate change. The current study demonstrates that N_2_O is the end reduction product when NO_3_^-^ is available, and N_2_O formation surpasses its consumption in the Siberian permafrost soils. Leveraging metagenomic analysis, the studies also reveal that both N_2_O formation and consumption are driven by non-denitrifying bacteria. While the detailed understanding of N_2_O formation and consumption has been well studied in non-permafrost soil and aquatic systems [19, 87], very limited information is available about the taxonomy, physiology, and ecology of the microorganisms involved in these two processes in permafrost. Further laboratory and field studies are needed to uncover the fate of the gaseous N species in rapidly changing high Arctic ecosystems, as well as to characterize microorganisms involved into N_2_O formation and consumption, potentially leading to more holistic prediction of N_2_O emissions in permafrost regions.

## Acknowledgments

The authors acknowledge funding through the NSF Dimensions of Biodiversity program (DEB-1442262 to T.A.V. and K.G.L. and DEB-1831599 to F.E.L), and the Genomic Science Program of the U.S. Department of Energy, Office of Science, Office of Biological and Environmental Research (DE-SC0020369 to K.G.L., and T.A.V.). Aspects of this work were carried out under the state assignment of the Institute of Physicochemical and Biological Problems of Soil Science, Russian Federation, Russian Academy of Sciences (RAS).

## Contributions

Y.S., X.W., F.E.L., and T.A.V. conceptualized the research and designed experiments. T.A.V., O.G.Z., E.M.R., and K.G.L. collected permafrost soil samples. Y.S. performed microcosm experiments, analytical work, DNA extraction, and bioinformatic analyses with help from X.W. Y.S. wrote the manuscript with input from all coauthors.

## Data availability

All metagenomic data sets were deposited in the European Nucleotide Archive under project PRJNA925153, and their respective accession numbers can be found in the Table S2.

## References

1. Gruber S. Derivation and analysis of a high-resolution estimate of global permafrost zonation. The Cryosphere. 2012;6:221–233.

2. Batjes NH. Total carbon and nitrogen in the soils of the world. Eur J Soil Sci. 1996;47:151–163.

3. Harden JW, Koven CD, Ping CL, Hugelius G, David McGuire A, Camill P, et al. Field information links permafrost carbon to physical vulnerabilities of thawing. Geophys Res Lett. 2012;39.

4. Hugelius G, Strauss J, Zubrzycki S, Harden JW, Schuur E, Ping C-L, et al. Estimated stocks of circumpolar permafrost carbon with quantified uncertainty ranges and identified data gaps. Biogeosciences. 2014;11:6573–6593.

5. Drake TW, Wickland KP, Spencer RG, McKnight DM, Striegl RG. Ancient low–molecular-weight organic acids in permafrost fuel rapid carbon dioxide production upon thaw. Proc Natl Acad Sci USA. 2015;112:13946–13951.

6. Voigt C, Marushchak ME, Lamprecht RE, Jackowicz-Korczyński M, Lindgren A, Mastepanov M, et al. Increased nitrous oxide emissions from Arctic peatlands after permafrost thaw. Proc Natl Acad Sci USA. 2017;114:6238–6243.

7. Voigt C, Marushchak ME, Abbott BW, Biasi C, Elberling B, Siciliano SD, et al. Nitrous oxide emissions from permafrost-affected soils. Nat Rev Earth Environ. 2020;1:420–434.

8. Lacroix F, Zaehle S, Caldararu S, Schaller J, Stimmler P, Holl D, et al. Mismatch of N release from the permafrost and vegetative uptake opens pathways of increasing nitrous oxide emissions in the high Arctic. Glob Change Biol. 2022;28:5973–5990.

9. Schuur EA, Vogel JG, Crummer KG, Lee H, Sickman JO, Osterkamp T. The effect of permafrost thaw on old carbon release and net carbon exchange from tundra. Nature. 2009;459:556–559.

10. Knoblauch C, Beer C, Liebner S, Grigoriev MN, Pfeiffer E-M. Methane production as key to the greenhouse gas budget of thawing permafrost. Nat Clim Change. 2018;8:309–312.

11. Christensen JH, Kanikicharla KK, Aldrian E, An SI, Cavalcanti IFA, de Castro M, et al. Climate phenomena and their relevance for future regional climate change. Cambridge University Press, 2013.

12. Lawrence DM, Slater AG, Swenson SC. Simulation of present-day and future permafrost and seasonally frozen ground conditions in CCSM4. J Clim. 2012;25:2207–2225.

13. Yin Y, Kara-Murdoch F, Murdoch RW, Yan J, Chen G, Xie Y, et al. Nitrous oxide inhibition of methanogenesis represents an underappreciated greenhouse gas emission feedback. ISME J. 2024;18:wrae027.

14. Marushchak M, Kerttula J, Diáková K, Faguet A, Gil J, Grosse G, et al. Thawing Yedoma permafrost is a neglected nitrous oxide source. Nat Commun. 2021;12:1–10.

15. IPCC. Climate change 1995: The science of climate change: contribution of working group I to the second assessment report of the Intergovernmental Panel on Climate Change, 1996.

16. Ravishankara A, Daniel JS, Portmann RW. Nitrous oxide (N_2_O): the dominant ozone- depleting substance emitted in the 21^st^ century. Science. 2009;326:123–125.

17. Etminan M, Myhre G, Highwood E, Shine K. Radiative forcing of carbon dioxide, methane, and nitrous oxide: a significant revision of the methane radiative forcing. Geophys Res Lett. 2016;43:12,614–12,623.

18. Elberling B, Christiansen HH, Hansen BU. High nitrous oxide production from thawing permafrost. Nat Geosci. 2010;3:332–335.

19. Knowles R. Denitrification. Microbiol Rev. 1982;46:43–70.

20. Wrage N, Velthof G, Van Beusichem M, Oenema O. Role of nitrifier denitrification in the production of nitrous oxide. Soil Biol Biochem. 2001;33:1723–1732.

21. Tiedje JM, Sexstone AJ, Myrold DD, Robinson JA. Denitrification: ecological niches, competition and survival. Antonie Van Leeuwenhoek. 1983;48:569–583.

22. Chalk P, Smith C. Chemodenitrification. Springer, 1983.

23. Onley JR, Ahsan S, Sanford RA, Löffler FE. Denitrification by *Anaeromyxobacter dehalogenans*, a common soil bacterium lacking the nitrite reductase genes *nirS* and *nirK*. Appl Environ Microbiol. 2018;84.

24. Sanford RA, Wagner DD, Wu Q, Chee-Sanford JC, Thomas SH, Cruz-García C, et al. Unexpected nondenitrifier nitrous oxide reductase gene diversity and abundance in soils. Proc Natl Acad Sci USA. 2012;109:19709–19714.

25. Abramov A, Vishnivetskaya T, Rivkina E. Are permafrost microorganisms as old as permafrost? FEMS Microbiol Ecol. 2021;97:fiaa260.

26. Monteux S, Keuper F, Fontaine S, Gavazov K, Hallin S, Juhanson J, et al. Carbon and nitrogen cycling in Yedoma permafrost controlled by microbial functional limitations. Nat Geosci. 2020;13:794–798.

27. Tecon R, Or D. Biophysical processes supporting the diversity of microbial life in soil. FEMS Microbiol Rev. 2017;41:599–623.

28. Liang R, Li Z, Lau Vetter MC, Vishnivetskaya TA, Zanina OG, Lloyd KG, et al. Genomic reconstruction of fossil and living microorganisms in ancient Siberian permafrost. Microbiome. 2021;9:1–20.

29. Wu X, Almatari AL, Cyr WA, Williams DE, Pfiffner SM, Rivkina EM, et al. Microbial life in 25-m-deep boreholes in ancient permafrost illuminated by metagenomics. Environ Microbiomes. 2023;18:1–19.

30. Rachlewicz G, Szczuciński W. Changes in thermal structure of permafrost active layer in a dry polar climate, Petuniabukta, Svalbard. Polish Polar Research. 2008;29:261–278.

31. Liang R, Lau M, Vishnivetskaya T, Lloyd KG, Wang W, Wiggins J, et al. Predominance of anaerobic, spore-forming bacteria in metabolically active microbial communities from ancient Siberian permafrost. Appl Environ Microbiol. 2019;85:e00560–00519.

32. Janssen H, Bock E. Profiles of ammonium, nitrite and nitrate in the permafrost soils. Viable microorganisms in permafrost Pushchino Research Centre Russian Academy of Sciences, Pushchino. 1994:27–36.

33. Löffler FE, Sanford RA, Ritalahti KM. Enrichment, cultivation, and detection of reductively dechlorinating bacteria. Meth Enzymol. 2005;397:77–111.

34. Sun Y, Yin Y, He G, Cha G, Ayala-del-Río HL, González G, et al. pH selects for distinct N_2_O-reducing microbiomes in tropical soil microcosms. ISME Commun. 2024:ycae070.

35. Wolin E, Wolin MJ, Wolfe R. Formation of methane by bacterial extracts. J Biol Chem. 1963;238:2882–2886.

36. Yin Y, Yan J, Chen G, Murdoch FK, Pfisterer N, Löffler FE. Nitrous oxide is a potent inhibitor of bacterial reductive dechlorination. Environ Sci Technol. 2018;53:692–701.

37. Zhang L, Yin Y, Sun Y, Liang X, Graham DE, Pierce EM, et al. Inhibition of methylmercury and methane formation by nitrous oxide in Arctic tundra soil microcosms. Environ Sci Technol. 2023;57:5655–5665.

38. Sander R. Compilation of Henry’s law constants for inorganic and organic species of potential importance in environmental chemistry. Max-Planck Institute of Chemistry, Air Chemistry Department Mainz, Germany, 1999. p^pp.

39. Bolger AM, Lohse M, Usadel B. Trimmomatic: a flexible trimmer for Illumina sequence data. Bioinformatics. 2014;30:2114–2120.

40. Li D, Liu C-M, Luo R, Sadakane K, Lam T-W. MEGAHIT: an ultra-fast single-node solution for large and complex metagenomics assembly via succinct de Bruijn graph. Bioinformatics. 2015;31:1674–1676.

41. Wu Y-W, Simmons BA, Singer SW. MaxBin 2.0: an automated binning algorithm to recover genomes from multiple metagenomic datasets. Bioinformatics. 2016;32:605–607.

42. Kang DD, Li F, Kirton E, Thomas A, Egan R, An H, et al. MetaBAT 2: an adaptive binning algorithm for robust and efficient genome reconstruction from metagenome assemblies. PeerJ. 2019;7:e7359.

43. Alneberg J, Bjarnason BS, De Bruijn I, Schirmer M, Quick J, Ijaz UZ, et al. Binning metagenomic contigs by coverage and composition. Nat Methods. 2014;11:1144–1146.

44. Uritskiy GV, DiRuggiero J, Taylor J. MetaWRAP—a flexible pipeline for genome-resolved metagenomic data analysis. Microbiome. 2018;6:1–13.

45. Parks DH, Imelfort M, Skennerton CT, Hugenholtz P, Tyson GW. CheckM: assessing the quality of microbial genomes recovered from isolates, single cells, and metagenomes. Genome Res. 2015;25:1043–1055.

46. Pasolli E, Asnicar F, Manara S, Zolfo M, Karcher N, Armanini F, et al. Extensive unexplored human microbiome diversity revealed by over 150,000 genomes from metagenomes spanning age, geography, and lifestyle. Cell. 2019;176:649–662. e620.

47. Boyd JA, Woodcroft BJ, Tyson GW. GraftM: a tool for scalable, phylogenetically informed classification of genes within metagenomes. Nucleic Acids Res. 2018;46:e59–e59.

48. McDonald D, Price MN, Goodrich J, Nawrocki EP, DeSantis TZ, Probst A, et al. An improved Greengenes taxonomy with explicit ranks for ecological and evolutionary analyses of bacteria and archaea. ISME J. 2012;6:610–618.

49. Chaumeil P-A, Mussig AJ, Hugenholtz P, Parks DH. GTDB-Tk: a toolkit to classify genomes with the Genome Taxonomy Database. Oxford University Press, 2020. p^pp.

50. Parks DH, Chuvochina M, Chaumeil P-A, Rinke C, Mussig AJ, Hugenholtz P. A complete domain-to-species taxonomy for Bacteria and Archaea. Nat Biotechnol. 2020;38:1079–1086.

51. Hyatt D, Chen G-L, LoCascio PF, Land ML, Larimer FW, Hauser LJ. Prodigal: prokaryotic gene recognition and translation initiation site identification. BMC Bioinform. 2010;11:1–11.

52. Aramaki T, Blanc-Mathieu R, Endo H, Ohkubo K, Kanehisa M, Goto S, et al. KofamKOALA: KEGG Ortholog assignment based on profile HMM and adaptive score threshold. Bioinformatics (Oxford, England). 2020;36:2251–2252.

53. Graham E, Heidelberg J, Tully B. Potential for primary productivity in a globally-distributed bacterial phototroph. ISME J. 2018;12:1861–1866.

54. Price MN, Dehal PS, Arkin AP. FastTree 2–approximately maximum-likelihood trees for large alignments. PLoS One. 2010;5:e9490.

55. Letunic I, Bork P. Interactive Tree Of Life (iTOL) v5: an online tool for phylogenetic tree display and annotation. Nucleic Acids Res. 2021;49:W293–W296.

56. R Core Team R. R: A language and environment for statistical computing. 2021.

57. Gómez-Rubio V. ggplot2-elegant graphics for data analysis. J Stat Softw. 2017;77:1–3.

58. McMurdie PJ, Holmes S. phyloseq: an R package for reproducible interactive analysis and graphics of microbiome census data. PLoS One. 2013;8:e61217.

59. Oksanen J, Blanchet FG, Kindt R, Legendre P, Minchin PR, O’hara R, et al. Package ‘vegan’. Community ecology package, version. 2013;2:1–295.

60. Kolde R, Kolde MR. Package ‘pheatmap’. R package. 2015;1:790.

61. Sullivan MJ, Gates AJ, Appia-Ayme C, Rowley G, Richardson DJ. Copper control of bacterial nitrous oxide emission and its impact on vitamin B_12_-dependent metabolism. Proc Natl Acad Sci USA. 2013;110:19926–19931.

62. Rustad L, Campbell J, Marion G, Norby R, Mitchell M, Hartley A, et al. A meta-analysis of the response of soil respiration, net nitrogen mineralization, and aboveground plant growth to experimental ecosystem warming. Oecologia. 2001;126:543–562.

63. Schaeffer SM, Sharp E, Schimel JP, Welker JM. Soil–plant N processes in a high Arctic ecosystem, NW Greenland are altered by long-term experimental warming and higher rainfall. Glob Change Biol. 2013;19:3529–3539.

64. Smith SL, O’Neill HB, Isaksen K, Noetzli J, Romanovsky VE. The changing thermal state of permafrost. Nat Rev Earth Environ. 2022;3:10–23.

65. Yang G, Peng Y, Marushchak ME, Chen Y, Wang G, Li F, et al. Magnitude and pathways of increased nitrous oxide emissions from uplands following permafrost thaw. Environ Sci Technol. 2018;52:9162–9169.

66. Overduin P, Portnov A, Ruppel C. NOAA Arctic report card 2023: permafrost beneath Arctic ocean margins. 2023.

67. Marushchak M, Pitkämäki A, Koponen H, Biasi C, Seppälä M, Martikainen P. Hot spots for nitrous oxide emissions found in different types of permafrost peatlands. Glob Change Biol. 2011;17:2601–2614.

68. Gil J, Pérez T, Boering K, Martikainen P, Biasi C. Mechanisms responsible for high N_2_O emissions from subarctic permafrost peatlands studied via stable isotope techniques. Global Biogeochem Cycles. 2017;31:172–189.

69. Hallin S, Philippot L, Löffler FE, Sanford RA, Jones CM. Genomics and ecology of novel N_2_O-reducing microorganisms. Trends Microbiol. 2018;26:43–55.

70. Shan J, Sanford RA, Chee-Sanford J, Ooi SK, Löffler FE, Konstantinidis KT, et al. Beyond denitrification: The role of microbial diversity in controlling nitrous oxide reduction and soil nitrous oxide emissions. Glob Chang Biol. 2021;27:2669–2683.

71. Meziti A, Rodriguez-R LM, Hatt JK, Peña-Gonzalez A, Levy K, Konstantinidis KT. The reliability of metagenome-assembled genomes (MAGs) in representing natural populations: insights from comparing MAGs against isolate genomes derived from the same fecal sample. Appl Environ Microbiol. 2021;87:e02593–02520.

72. Jones CM, Graf DR, Bru D, Philippot L, Hallin S. The unaccounted yet abundant nitrous oxide-reducing microbial community: a potential nitrous oxide sink. ISME J. 2013;7:417–426.

73. Zhou J, Ning D. Stochastic community assembly: does it matter in microbial ecology? Microbiol Mol Biol R. 2017;81:10.1128/mmbr.00002-00017.

74. Gilichinsky D, Wagener S, Vishnevetskaya T. Permafrost microbiology. Permafr Periglac Process. 1995;6:281–291.

75. Vorobyova E, Soina V, Gorlenko M, Minkovskaya N, Zalinova N, Mamukelashvili A, et al. The deep cold biosphere: facts and hypothesis. FEMS Microbiol Rev. 1997;20:277–290.

76. Khlebnikova G, Gilichinskii D, Fedorov-Davydov D, Vorob’eva E. Quantitative evaluation of microorganisms in permafrost deposits and buried soils. Microbiology (New York). 1990;59:106–112.

77. Mariangel L, Aspe E, Cristina Marti M, Roeckel M. The effect of sodium chloride on the denitrification of saline fishery wastewaters. Environ Technol. 2008;29:871–879.

78. Vishnivetskaya T, Kathariou S, McGrath J, Gilichinsky D, Tiedje JM. Low-temperature recovery strategies for the isolation of bacteria from ancient permafrost sediments. Extremophiles. 2000;4:165–173.

79. Mackelprang R, Burkert A, Haw M, Mahendrarajah T, Conaway CH, Douglas TA, et al. Microbial survival strategies in ancient permafrost: insights from metagenomics. ISME J. 2017;11:2305–2318.

80. Glass C, Silverstein J. Denitrification of high-nitrate, high-salinity wastewater. Water Res. 1999;33:223–229.

81. Huang J, Han M, Yang J, Kappler A, Jiang H. Salinity impact on composition and activity of nitrate-reducing Fe (II)-oxidizing microorganisms in saline lakes. Appl Environ Microbiol. 2022;88:e00132–00122.

82. Nikrad MP, Kerkhof LJ, Häggblom MM. The subzero microbiome: microbial activity in frozen and thawing soils. FEMS Microbiol Ecol. 2016;92:fiw081.

83. Feller G, Gerday C. Psychrophilic enzymes: hot topics in cold adaptation. Nat Rev Microbiol. 2003;1:200–208.

84. Pörtner H-O, Roberts DC, Masson-Delmotte V, Zhai P, Tignor M, Poloczanska E, et al. The ocean and cryosphere in a changing climate. IPCC special report on the ocean and cryosphere in a changing climate. 2019;1155.

85. McGuire AD, Lawrence DM, Koven C, Clein JS, Burke E, Chen G, et al. Dependence of the evolution of carbon dynamics in the northern permafrost region on the trajectory of climate change. Proc Natl Acad Sci USA. 2018;115:3882–3887.

86. Biskaborn BK, Smith SL, Noetzli J, Matthes H, Vieira G, Streletskiy DA, et al. Permafrost is warming at a global scale. Nat Commun. 2019;10:264.

87. Zumft WG. Cell biology and molecular basis of denitrification. Microbiol Mol Biol R. 1997;61:533–616.

